# Peripheral mitochondrial function correlates with clinical severity in idiopathic Parkinson’s disease

**DOI:** 10.1101/422089

**Authors:** Chiara Milanese, Cesar Payan-Gomez, Marta Galvani, Nicolás Molano González, Maria Tresini, Soraya Nait Abdellah, Willeke M.C. van Roon-Mom, Silvia Figini, Johan Marinus, Jacobus J. van Hilten, Pier G. Mastroberardino

## Abstract

**Background:** Parkinson disease is an intractable disorder with heterogeneous clinical presentation that may reflect different underlying pathogenic mechanisms. Surrogate indicators of pathogenic processes correlating with clinical measures may assist in better patients stratification. Mitochondrial function - which is impaired in and central to PD pathogenesis - may represent one of such surrogate indicators.

**Methods:** Mitochondrial function was assessed by respirometry experiment in fibroblasts derived from idiopathic patients (n=47) in normal conditions and in experimental settings that do not permit glycolysis and therefore force energy production through mitochondrial function. Respiratory parameters and clinical measures were correlated with bivariate analysis. Machine learning based classification and regression trees were used to classify patients on the basis of biochemical and clinical measures. Effects of mitochondrial respiration on alpha-synuclein stress was assessed monitoring the protein phosphorylation in permitting versus restrictive glycolysis conditions.

**Results:** Bioenergetics properties in peripheral fibroblasts correlate with clinical measures in idiopathic patients and correlation is stronger with predominantly non-dopaminergic signs. Bioenergetics analysis under metabolic stress, in which energy is produced solely by mitochondria, shows that patients’ fibroblasts can augment respiration, therefore indicating that mitochondrial defects are reversible. Forcing energy production through mitochondria, however, favors alpha-synuclein stress in different cellular experimental systems. Machine learning-based classification identified different groups of patients in which increasing disease severity parallels higher mitochondrial respiration.

**Conclusion:** Suppression of mitochondrial activity in Parkinson disease may be an adaptive strategy to cope with concomitant pathogenic factors. Moreover, mitochondrial measures are potential biomarkers to follow disease progression.

## Introduction

Parkinson disease (PD) is a multisystem disorder characterized by a broad spectrum of motor (stiffness, slowness, tremor, gait and balance difficulties) and non-motor (cognitive, psychiatric, sleep, alertness, autonomic) disturbances. The latter may antedate the motor symptoms, worsen as the disease advances, and predominate in causing disability during the later stages of the disease ^1^. PD patients do not follow a uniform disease course, but exhibit conspicuous differences in primary disease-related and medication-induced complications as well as in the rate of progression of the disease, reflecting the existence of subtypes. In early PD, however, clinical characteristics are insufficient to identify subtypes ^2^. There is, however, consensus that age at onset of manifest disease is a major determinant of progression, with a more progressive course being associated with a higher age at onset ^3^.

Better patient stratification is essential in designing clinical trials and may be achieved by integrating clinical data with quantitative biomarkers to best reflect the progression of the disease and its underlying biological pathophysiology. Furthermore, these biomarkers may have the potential to identify systems or persons at-risk before overt expression of the disorder and will allow earlier diagnosis and faster evaluation of clinical trials outcome. Ideal surrogate measures reflect processes in a crucial causal pathway of the disease and correlate with the true clinical outcome ^4^.

Dysfunction in mitochondrial oxidative phosphorylation (OXPHOS) has been linked to PD by multiple sources of converging evidence including genetic, toxicological, and epidemiological studies ^5-7^. Mitochondrial complex I damage induces parkinsonism in humans and models the diseases in laboratory animals; moreover, mitochondrial defects are detectable in peripheral cells of genetic and idiopathic PD cases ^6,8-10^. In addition, interventions targeting mitochondria ameliorate pathology in multiple animal models and improve respiration efficiency in patients’ fibroblasts ^11^. Collectively, these elements indicate that mitochondrial parameters might serve as an alternative outcome to complement clinical measures.

Here we performed bioenergetic characterization in primary fibroblasts from a highly characterized cohort of 47 PD patients to test the hypothesis that mitochondrial parameters correlate with clinical features and may therefore be informative of the clinical outcome. We applied statistical models and machine learning procedures to describe the complex relationship between the different analyzed parameters and achieved unbiased grouping of patients on the basis of both clinical and laboratory measures. To fully expose mitochondrial defects, we performed biochemical experiments also under conditions of metabolic stress where the function of glycolysis - which could compensate and hide mitochondrial anomalies – is minimized. Finally, we explored detrimental synergies between bioenergetics and alphs-synuclein pathology in primary fibroblasts and differentiated neurons.

## Materials and Methods

### Patients

The present cross-sectional study in PD patients is part of the PROPARK (PROfiling PARKinson’s disease) study. Patients were recruited from the outpatient clinic for Movement Disorders of the Department of Neurology of the Leiden University Medical Center (LUMC; Leiden, the Netherlands) and nearby university and regional hospitals. All participants fulfilled the United Kingdom Parkinson’s Disease Society Brain Bank criteria for idiopathic PD ^12^. Evaluations occurred between January 2013 and January 2016. Exclusion criteria were: previous or other disorders of the central nervous system, peripheral nerve disorders influencing motor and/or autonomic functioning, and psychiatric comorbidity not related to PD.

All patients, except for 18 dopaminergic drug-naïve patients, were tested while on dopaminergic medication. The severity of motor symptoms was quantified using the Movement Disorder Society version of the Unified Parkinson’s Disease Rating Scale (MDS-UPDRS) motor examination (part III) ^13^. Additionally, the SENS-PD scale was administered, which is a composite score comprising three items with four response options (0-3) from each of the following six domains: postural instability and gait difficulty, psychotic symptoms, excessive daytime sleepiness, autonomic dysfunction, cognitive impairment and depressive symptoms (total range: 0-54) ^1^. These six domains represent a coherent complex of symptoms that largely do not improve with dopaminergic medication that is already present in the early disease stages and increases in severity when the disease advances ^14^. Higher scores on both scales reflect more severe impairment. Cognitive performance was assessed using the SCOPA-COG (SCales for Outcomes in PArkinson’s disease-COGnition; range 0-43), a valid and reliable instrument examining the following domains: memory, attention, executive functioning and visuospatial functioning ^15^; lower scores reflect more severe impairment. A levodopa dose equivalent (LDE) of daily levodopa (LDE-DOPA), dopamine agonists (LDE-DA), as well as a total LDE was calculated according to the formula developed by Tomlinson et al. ^16^.

The study was approved by the medical ethics committee of the LUMC and written informed consent was obtained from all PD patients.

### Fibroblasts cultures

PD patients’ fibroblasts were prepared isolated at LUMC from skin biopsies derived from the ventral side of the upper leg and cultured under highly standardized conditions at 37°C and 5% CO_2_ up to a maximum of 10 passages. Number of passages was kept consistent within groups. Fibroblasts were cultured in glucose (glucose 10 mM, 10% FBS (F6178, Sigma-Aldrich), 2 mM glutamine, 5 mM Hepes and 1% penicillin-streptomycin (P4333, Sigma-Aldrich)) or galactose (galactose 10 mM, 10% FBS, 2 mM glutamine, 5 mM Hepes and 1% penicillin-streptomycin) medium. Fibroblasts from sex-matched controls of comparable age (identification codes AG04659, AG05266, AG06283, AG07936, AG08152, AG08268, AG08269, AG08543, AG09162, AG09271, AG09879, AG12428, AG12951, AG13077, AG13348) were obtained from the Coriell biorepository of the Coriell Institute for Medical Research (Camden, NJ, USA).

### SH-SY5Y cell culture

SH-SY5Y neuroblastoma cells were maintained in DMEM medium, 10% FCS and 1% pen/strep. Cells were seeded at a density of 10 × 10^4^ on coverslips coated with laminin (L4544, Sigma) and poly-Ornithine (P4957, Sigma), and were differentiated for 8 days in DMEM 1% FCS containing retinoic acid (10µM, R2625 Sigma).

### Mitochondrial respiration and glycolysis determination

Bioenergetic profiles of human primary skin fibroblasts were generated in real time with a Seahorse XF24 Extracellular Flux Analyzer (Agilent Technologies, Santa Clara, Ca, USA) as previously described ^8, 11^. Fibroblasts were seeded on a Seahorse XF-24 plate at a density of 6×10^4^ cells per well and grown overnight in DMEM (10% of FCS and 1% Pen-Strep) at 37° C, 5% CO_2_. This density ensures a proportional response to the uncoupler FCCP (Carbonyl cyanide 4-(trifluoromethoxy)phenylhydrazone) with cell number ^8,11^ and resulted in confluent cultures, in which cell replication was further prevented by contact inhibition. On the experimental day, medium was changed to unbuffered DMEM (XF Assay Medium – Agilent Technologies, Santa Clara, Ca, USA) supplemented with 5 mM glucose and 1 mM sodium pyruvate, and incubated 1 hour at 37° C in the absence of CO_2_. Medium and reagents acidity was adjusted to pH 7.4 on the day of the assay. After four baseline measurements for the oxygen consumption ratio (OCR), cells were sequentially challenged with injections of mitochondrial toxins: 0.5 µM oligomycin (ATP synthase inhibitor), 1 µM FCCP (mitochondrial respiration uncoupler), 0.5 µM rotenone (complex I inhibitor), and 0.5 µM antimycin (complex III inhibitor).

### Lentiviral-mediated alpha-synuclein infection and immunocytochemistry procedures

For lentivirus production, HEK 293T cells were transfected with the pLenti-DsRed-IRES-SNCA:EGFP expression vector (^17^, Adgene 92195), together with psPAX2 (Addgene 12260), and pMD2.G (Addgene 12259) packaging and envelope plasmids, at (1: 0.75: 0.25 ratios respectively) using FuGENE 6 (E2691, Promega) according to the manufacturer’s instructions. Viral supernatants were harvested 48 h post-transfection, filtered through 0.45 micron filters (Millipore Corp.) and used immediately to infect subconfluent cell cultures of primary human dermal fibroblast derived from healthy donors and PD patients in the presence of 4 ug/ml polybrene (Hexadimethrine bromide polybrene, H9268, Sigma).

Forty-eight hours after infection, virus was removed and cells were incubated with glucose or galactose-containing medium and then processed for immunocytochemistry. Briefly, cells were fixed in PFA 4% for 20’ and, after washes in PBS, blocked in 3% BSA, 0,1% Triton X-100 and PBS for additional 20’. Cells were then incubated with an anti-GFP antibody (1:1000; 14314500; Roche) recognizing the GFP-alpha-synuclein fusion protein and with the anti-phosphorylated alpha-synuclein (S129) antibody (1:1000; ab59264, Abcam) for 3 hours at room temperature (RT). After 3 washes with PBS, cells were incubate for 1 hour with secondary antibodies Alexa 488 conjugated donkey anti-mouse IgG 1:1000 (Invitrogen), Cy5-conjugated donkey anti-rabbit IgG 1:1000 (Jackson) and DAPI. Image acquisition was performed in a Zeiss SP5 confocal microscope. The detection parameters were set in the control samples and were kept constant across specimens. Images were analyzed in a semi-automated fashion using constant thresholding parameters with the Metamorph software (Molecular Devices).

### Statistical analysis

Statistical associations were determined using classical bivariate analysis: Kruskall-Wallis test was used for comparison of quantitative against categorical variables, Chi-square test was used for comparison of categorical vs. categorical variables, and Spearman correlation coefficient was used for comparison of quantitative vs. quantitative variables. The significance level was set at 0.05. Individual mitochondrial parameters were compared to the groups means using one-way ANOVA and Dunnett’s test. A modified t-test as described in Crawford et al. ^18^ provided conceptually comparable results (data not shown).

Stratification was achieved using applied classification and regression trees (CART) ^19^. The rpart package ^20^ in R software ^21^ was used to fit data into CART; the function rpart was used with ANOVA. All statistical analyses were performed in R version 3.3.2.

## Results

### Characterization of mitochondrial function in permitting *versus* non-permitting glycolysis conditions

#### Bioenergetics analysis

Initially, we characterized bioenergetics properties in PD patients’ fibroblasts (N=47) and sex-matched health subjects of comparable age (N=13). We used a Seahorse Extracellular flux analyzer because this instrument returns multiple bioenergetics parameters related to both mitochondrial function and glycolysis in a single experiment (fig. 1A). When analyzed in glucose-supplemented medium, where glycolysis is permitted, PD pooled data did not show differences in basal respiration, reserve capacity, or rotenone sensitive respiration (i.e. complex I driven) when compared to the respective controls (fig. 1B). To unravel defects that are eventually masked in glycolysis permitting conditions, we repeated the experiments replacing glucose with galactose, which forces cells to rely on mitochondria for ATP production. Of note, these culturing conditions are lethal for fibroblasts from patients with mitochondrial pathologies ^22^. PD patients’ fibroblasts, however, perfectly survived in galactose medium and no cell death was observed (data not shown); moreover, pooled respiration data revealed differences only in the reserve capacity, which was slightly reduced in PD patients (fig. 1C).

**Figure 1.**
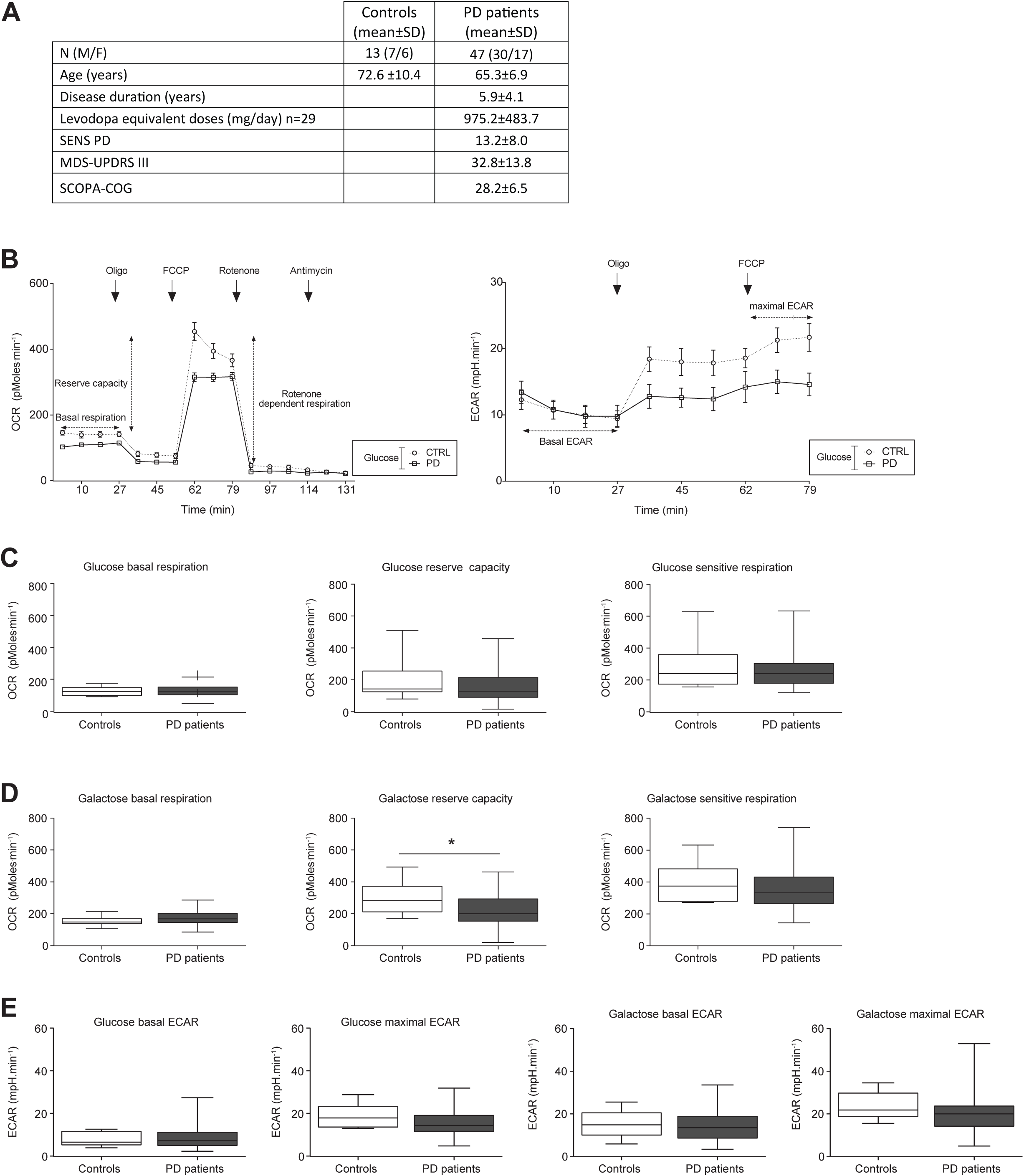
(**A**) Clinical description of cases and controls included in this research. (**B**) Representative Seahorse traces of oxygen consumption rate (OCR, left) and extracellular acidification rate (ECAR, right) reflecting mitochondrial respiration and glycolysis respectively. Physiological measures occur in real time with sequential injection of oligomycin (ATP synthase inhibitor), FCCP (uncoupler), rotenone (complex I inhibitor), and antimycin (complex III inhibitor). (**C**) Respiratory parameters of pooled bioenergetics raw data in glucose medium, i.e. in glycolysis permitting conditions. (**D**) Respiratory parameters of pooled bioenergetics raw data in galactose medium. (**E**) ECAR is not different in all the tested experimental conditions. * p<0.0332, Kruskal-Wallis non-parametric test.

In agreement with previous analysis on idiopathic PD ^8^ or Parkin mutant fibroblasts ^23^, we did not observe any difference in basal or stimulated extracellular acidification rate (ECAR) in both glucose or galactose culturing conditions (Fig 1D).

While analysis on pooled data failed to reveal major differences between PD and control groups, when analyzed at individual level – i.e. single patients where compared to the average of the control group – the results exposed significant variability in both glucose and galactose cultured specimens (fig. 2A, B). Forcing bioenergetics through oxidative metabolism (i.e. galactose medium) unmasked anomalies and several lines that did not exhibit alterations in glucose medium revealed differences in basal respiration when cultured with galactose (fig. 2C, D). However, no changes were detected in reserve capacity and rotenone sensitive respiration (fig. 2D).

**Figure 2.**
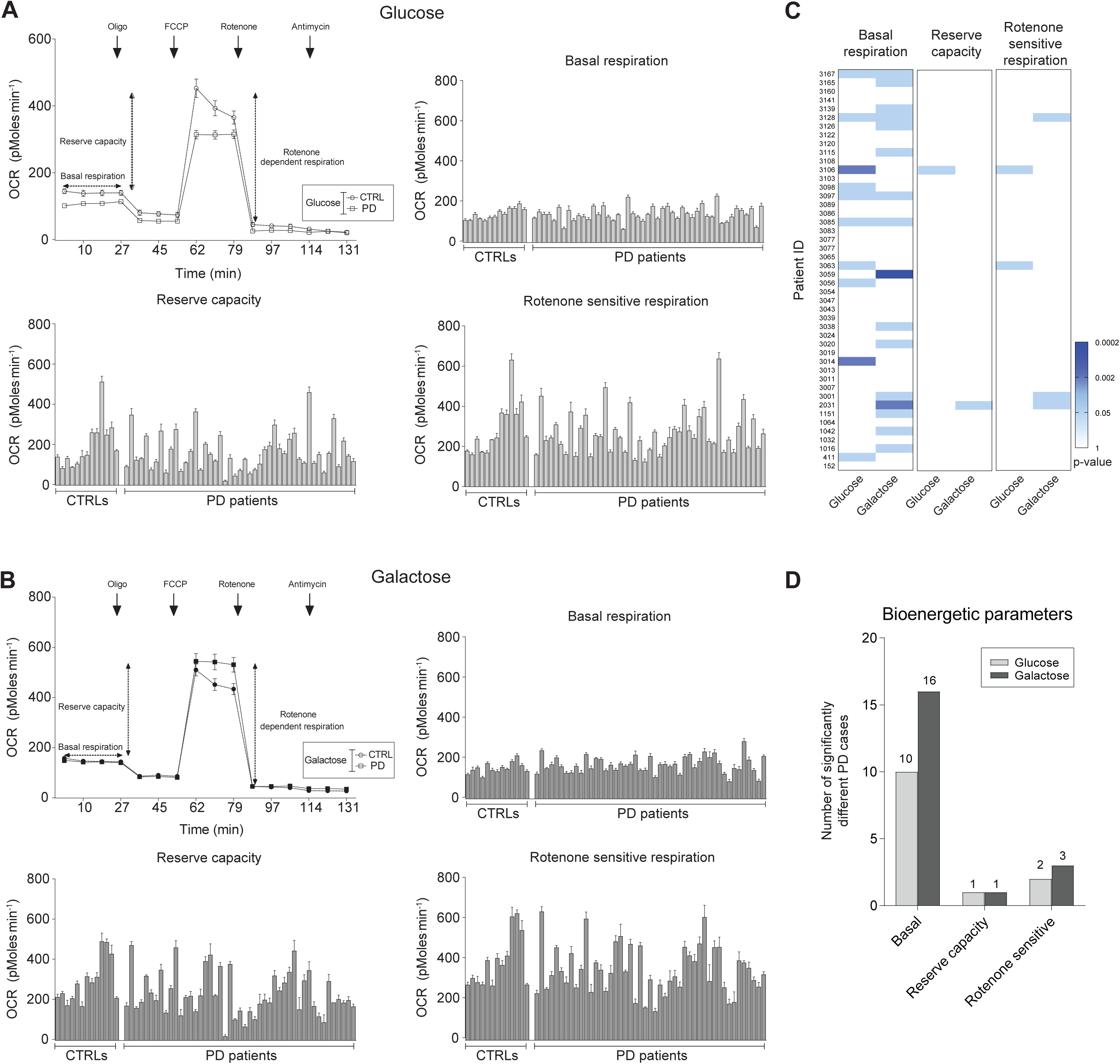
Analysis of individual bioenergetics data highlights high variability in reserve capacity and in rotenone sensitive respiration in both glucose (**A**) and galactose (**B**) medium conditions. (**C**) Heat map plotting statistical significance values of the differences between individual patients’ respiration data and the mean of healthy controls. Significance was determined using one-way ANOVA and Dunnett’s test. (**D**) Bar graph showing the number of patients with statistically significant difference in respiratory parameters.

#### PD fibroblasts can augment respiration in conditions forcing metabolism through oxidative phosphorylation

The adaptation of mitochondrial function to conditions that force bioenergetics through oxidative metabolism (i.e. galactose medium) is reflected in the ratio between respiration parameters obtained in galactose and glucose conditions (galactose-to-glucose ratio).

In cells from healthy controls, galactose medium augmented respiration (i.e. galactose-to-glucose ratio>1) in most lines (10 out of 13) and only three lines displayed suppression in more than two of the analyzed respiratory parameters (fig. 3A, B).

**Figure 3.**
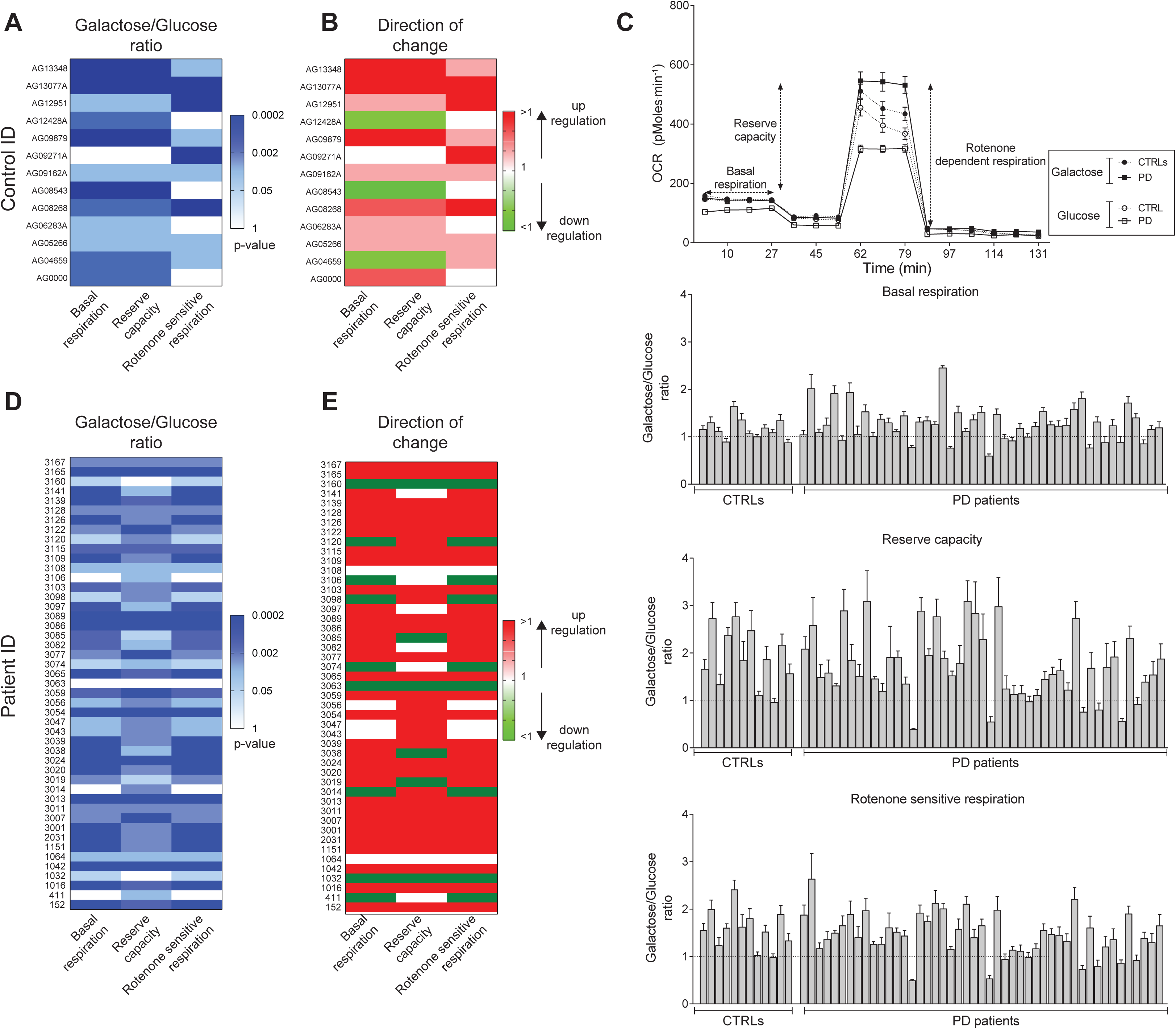
Analysis of the galactose-to-glucose respiration ratio as an index of mitochondria alterations induced by conditions non permitting glycolysis. (**A**) Replacing glucose with galactose alters respiration parameters in control cells. The heat map illustrates statistical significance values obtained testing the hypothesis that in control specimens the galactose-to-glucose respiration ratios are different than one, which would indicate altered respiration in non-permitting versus permitting glycolysis conditions. (**B**) Heat map illustrating the direction of the changes in the galactose-to-glucose ratios; red indicates a ratio higher than one, i.e. upregulation of respiration in galactose. The vast majority of control lines potentiates respiration when bioenergetics is forced through OXPHOS. (**C**) Representative Seahorse trace and histograms of respiratory parameters of individual bioenergetics data expressed as galactose over glucose ratio. (**D**) Heat map displaying statistical significance values obtained testing the hypothesis that in PD specimens the galactose-to-glucose respiration ratios are different than one. (**E**) Heat map illustrating the direction of the changes in the galactose-to-glucose ratios; red indicates a ratio higher than one, i.e. upregulation of respiration in galactose. The vast majority of PD lines is able to augment respiration when bioenergetics is forced through OXPHOS. Significance was determined using one-way ANOVA and Dunnett’s tests.

Galactose medium altered mitochondrial function also in PD fibroblasts, albeit the response was variable among the different lines (fig. 3C, D). The direction of the changes observed in the patient specimens, however, was unexpected. In fact, given the mitochondrial defects intrinsic to PD, which are observed also peripherally, one would predict inability to augment respiration and even lethality – i.e. the opposite outcome than in control cells - as reported for typical mitochondrial disorders ^22^. However, basal respiration, reserve capacity, and rotenone sensitive respiration significantly increased in several PD fibroblast lines, while none of the analyzed PD specimens displayed decreased respiratory parameters (fig. 3E). This evidence indicates that, rather than being irreversibly compromised, mitochondrial dysfunction in PD can be restored under certain metabolic conditions.

### Correlation between laboratory and clinical measures

We next examined whether the variability observed in respiration parameters might reflect clinical characteristics.

No significant correlations (r_s_) were found between respirometry parameters and age, age of onset, disease duration or levodopa equivalent doses; in addition, the severity of motor symptoms did not correlate with any of the laboratory parameters (Fig. 4A). However, the SENS-PD scores correlated with glucose reserve capacity (r_s_ = .342, p = .026), while the SCOPA-COG score displayed significant correlations with reserve capacity in the glucose medium (r_s_ = −.370, p = .017) and rotenone sensitive respiration (r_s_ = −0.320, p=.041) in the glucose medium (fig. 4A, B).

**Figure 4.**
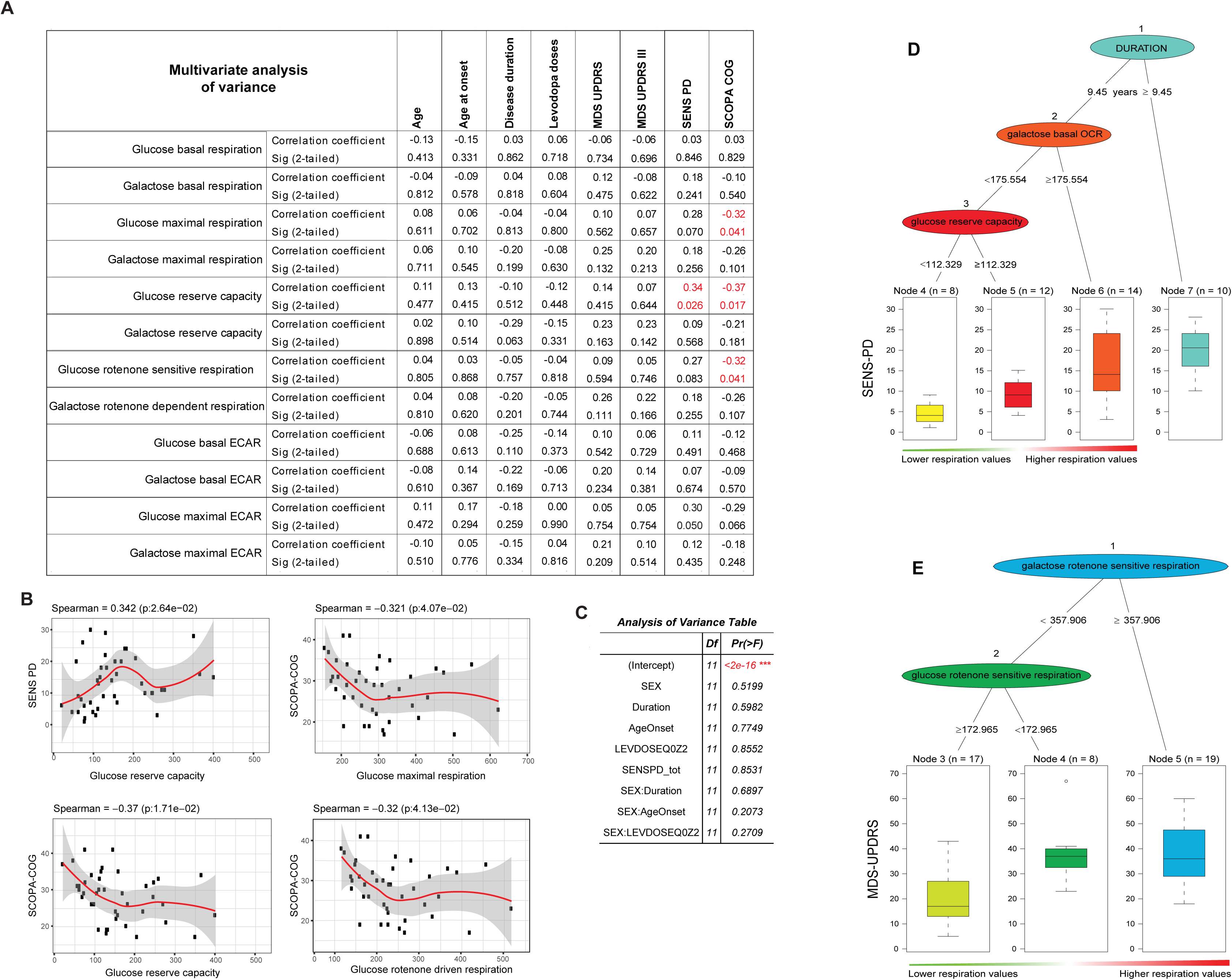
Correlation between raw respiration data and clinical measures. (**A**) Multivariate analysis of variance showing Spearman’s correlation coefficients between laboratory and clinical measures and related significance. (**B**) Graphs of clinical and raw laboratory variables displaying statistically significant correlations. (**C**) Linear regression with interactions and analysis of variance indicates that correlation between the clinical and laboratory measures is independent from gender, age, age of onset, duration of the disease, and medication. (**D**) Grouping of patients using unbiased classification and regression tree (CART) analysis using the SENS-PD as response variable. (**E**) CART analysis using the MDS-UPDRS score as response variable.

### Unbiased grouping of patients on the basis of laboratory and clinical measures

The heterogeneity in clinical presentation, the dispersion we observed in mitochondrial physiology, and their correlation lend support to the hypothesis that there may be sub-groups of different mitochondrial phenotypes correlating with the symptomatology in the general population of idiopathic PD patients. To test this possibility, we used machine learning methodology and used recursive partitioning to build a classification and regression tree (CART) ^24^, which have been successfully used in applications such as clinical subtypes classification ^25^ and neuroimaging data analysis to predict Alzheimer’s disease ^26^.

All available parameters – i.e. demographic variables (i.e. age, age of at onset, duration of the disease, and gender), the equivalent of levodopa medication, respirometry and acidification parameters – were used in the CART process as input variables to predict either the SENS-PD or the MDS-UPDRS III (i.e. response variables). Intrinsic to CART modeling is a selection step that eliminates in an unbiased fashion redundancy among the input variables to identify the most significant parameters.

When CART analysis was applied using SENS-PD as a response variable, three rules - i.e. nodes 1, 2, and 3 - grouped the cases in four classes (Fig. 4D). The first rule, node A, identifies disease duration as a classifying variable and identifies a class of patients (Class 4) with disease duration longer than 9.45 years presenting with the most severe symptoms. The second and third rules - node 2 and 3 - respectively identify basal respiration in galactose and reserve capacity in glucose medium as classifying variables and divide patients with disease duration shorter than 9.45 years in three further classes. Class 1 is defined by lower galactose basal respiration (<175.5 pmol·min^-1^), lower glucose reserve capacity (<112.5 pmol·min^-1^), and includes patients with the least severe clinical presentation. Class 2 includes patients with lower galactose basal respiration, but higher glucose reserve capacity, and in class 3 both respiration parameters are higher. Presentation in these two classes is more severe than in class 1, with class 3 encompassing patients with worse symptoms. Overall, these data indicate that in patients with shorter disease duration (i.e. less than 9.45 years) higher respiration is associated with increasing symptoms’ severity.

When the MDS-UPDRS was used as response variable, CART analysis returned only two rules (node 1, rotenone sensitive respiration in galactose, node 2, rotenone sensitive respiration in glucose) dividing the patients in three classes (fig. 4E). Also in this case, higher values in respiration parameters are associated with more severe symptoms.

### Increased mitochondrial function promotes α-syn stress *in vitro*

Taken together, our combined biochemical and clinical data indicate that mitochondrial function is suppressed in PD and – given that lower respiration correlates with less severe symptoms – this alteration may reflect a protective adaptation to counterbalance PD pathogenesis. As a corollary, high mitochondrial activity may synergize with other pathogenic factors to elicit deterioration. Given that α-synuclein (α-syn) aggregation and stress are hallmarks of PD, we hypothesized that forcing mitochondrial activity in galactose medium could favor synucleinopathy. Investigating these aspects in fibroblasts, however, poses critical issues because α-syn expression is extremely low in this cell type ^27^. We therefore took advantage of lentiviral technology to engineer fibroblasts to express GFP-tagged human α-syn in three control (AG08268, AG08543, and AG13077) lines and three PD lines able to upregulate mitochondrial function in galactose (3020, 3039, and 3086). We evaluated protein stress by measuring the number of foci of phosphorylated α-syn (p-syn), which is the principal modified species of a-syn within PD pathological inclusions ^28^; intracellular foci therefore reflect early forms of aggregation.

In control cells, galactose medium conditions caused significant increase in the number of p-syn *foci* – which were quantified by an unbiased semi-automated procedure - therefore indicating that increased mitochondrial function may indeed favor synucleinopathy. In PD fibroblasts p-syn levels were elevated also in glucose conditions and did not show any further increase in galactose (fig. 5A, B). These effects were not caused by different levels of α-syn-GFP because the exogenous protein was expressed at comparable levels in control and PD specimens (fig. 5C – green signal).

**Figure 5.**
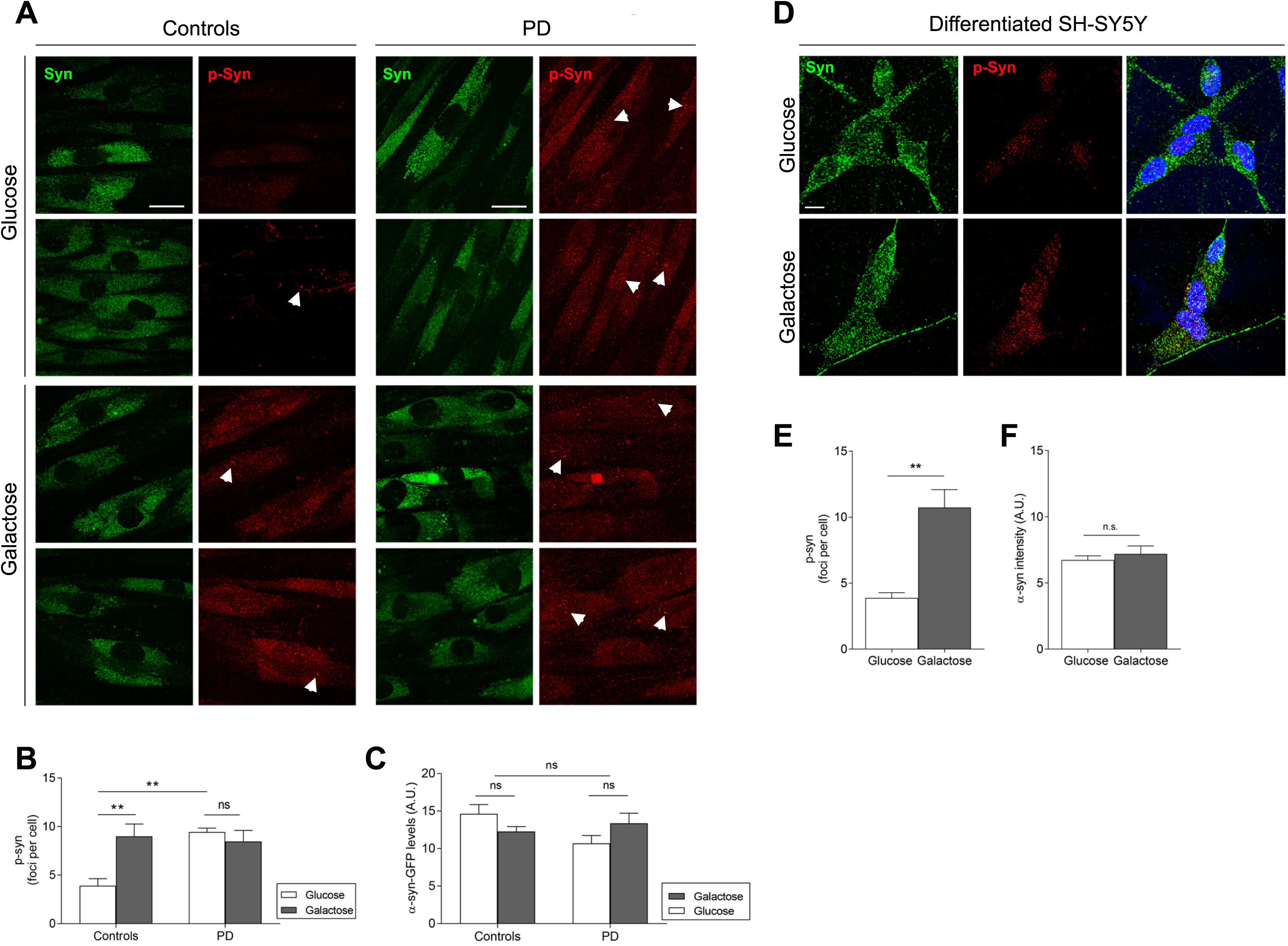
Increased mitochondrial function in galactose medium favors α-syn stress. (**A**) Representative laser scanning confocal microscopy imaging showing GFP-tagged α-syn (green) and p-syn (red) levels. In healthy controls (N=3), galactose significantly increases the number of intracellular p-syn *foci* (arrowheads) pointing to α-syn stress. In PD cells (N=3), p-syn levels are elevated also in glucose conditions and do not increase in galactose medium. (**B**) Quantification of intracellular p-syn *foci*. (**C**) Quantification of α-syn-GFP levels indicating comparable levels in control and PD specimens. (**E**) Representative laser scanning confocal microscopy imaging of differentiated SH-SY5Y cells showing endogenous α-syn (green) and p-syn (red) levels in glucose or galactose culturing conditions. (**F**) Quantification of intracellular p-syn foci showing increased α-syn stress in galactose medium. (**G**) Quantification of endogenous α-syn shows no differences between the two culturing conditions * p<0.0332, ** p<0.0021, Kruskal-Wallis non-parametric test. Scale bar = 10 µm.

To determine whether forcing bioenergetics metabolism through mitochondrial function aggravates endogenous α-syn stress also in neuronal cells, we investigated differentiated SH-SY5Y cells. This dopaminergic cell line expresses detectable levels of endogenous α-syn, which do not differ between glucose or galactose culturing conditions. However, cells grown in galactose medium exhibited significantly increased levels of p-syn, confirming the data obtained in overexpressing cells. Taken together, these findings indicate that the complex genetic background of idiopathic patients promotes per se α-syn stress and that suppression of mitochondrial function to mitigate synucleinopathy is ineffective in PD cells.

## Discussion

The data presented in this study are completely consistent and confirm previous observations from others and our laboratories showing impairment in mitochondrial function in PD ^6, 8^. However, in this study we report the unexpected and surprising finding that mitochondrial function in PD patients peripheral fibroblasts can be potentiated in conditions forcing OXPHOS, i.e. in galactosemedium. This evidence represents a paradigm shift in the current view of mitochondria in PD and suggests that, rather than being irreversibly damaged, mitochondrial function is suppressed. A possible hypothesis to explain this observation is that in PD mitochondria may suffer from intrinsic anomalies resulting in harmful consequences if the organelles function at normal levels and/or that mitochondrial activity may promote other PD-related pathogenic processes. Consistently with the latter hypothesis, galactose medium induced an increase in α-syn stress – indicated by increased p-syn levels – in GFP-syn expressing fibroblasts from healthy subjects. Experiments in engineered fibroblasts and differentiated SHSY-5Y cells also revealed that α-syn stress occurs at high levels in PD cells also when grown in glucose and does not increase in galactose, therefore indicating that the complex genetic background of idiopathic PD patients promotes α-syn stress *per se*. A protective role for suppression of mitochondrial function is consistent with recent hypotheses suggesting that neurodegeneration in Alzheimer’s disease is caused by metabolic alterations and that dysfunctional neurons upregulate mitochondrial respiration according to an inverse-Warburg effect pathogenic paradigm ^29,30^. Consistently, we have recently demonstrated that mild inhibition of complex I caused by nitrite-mediated complex I S-nitrosation is protective in multiple models of PD and improves bioenergetics efficiency in fibroblasts of PD patients, but not in matched controls ^11^. On these premises, our data suggest that – at least in some subtypes of PD – mitochondrial function is an amenable target for disease modification.

Two mitochondrial variables – reserve capacity and rotenone sensitive (i.e. complex I driven) respiration - correlate with the SENS-PD scale. These results confirm the pivotal role of complex I in PD pathobiology and are consistent with previous studies indicating that reserve capacity is very sensitive to stress and therefore particularly suited to detect systemic physiological anomalies ^31,32^. The SENS-PD scale addresses clinical features that mostly do not improve on dopaminergic treatment ^14^. It is likely that these predominantly non-dopaminergic items more accurately reflect severity and progression of the underlying disease pathobiology ^1^.

We used CART analysis ^19^ - a machine learning methodology using recursive partitioning - to classify patients on the basis of clinical and bioenergetic measures. CART - which have been already used to analyze biomedical problems ^33^ and have been used to stratify patients ^25^ - have several advantages over traditional approaches such as generalized linear models (e.g. linear regression, logistic regression among others). First, the method is non-parametric and thus does not assume any distribution model for the dependent variable. Second, it can handle a large set of explanatory variables and it automatically selects the most important variables to be used in the final model. Compared to classical regression methods where variable selection is an open problem with no definitive answer ^34^, CART analysis is data-driven and identifies interactions objectively, in an unbiased manner, and does not require any input from the researcher.

Not surprisingly, the highest hierarchical discriminant of clinical phenotypes is disease duration, with longer durations associated with more severe clinical presentation. The other classifying variables selected by CART among the multitude of demographic, clinical, and bioenergetics parameters are related to mitochondrial function and the analysis indicates that patients falling in classes with higher respiration present with increased severity of symptoms. These data substantiate the relevance of mitochondrial biology in PD pathogenesis and further support the hypothesis that suppression of mitochondrial function during PD pathogenesis might represent a protective adaptive response.

Using the SENS-PD scale as response variable led to better separation of patients with shorter disease duration and milder symptoms, therefore confirming the concept that signs outside the dopaminergic domain – which are less sensitive to dopaminergic medications, may manifest at earlier stages, and are enriched in the SENS-PD scale - can be highly informative for PD phenotyping and to monitor the disease progression ^1^.

In summary, our study reveals new aspects of mitochondrial biology in PD, establishes a connection between clinical and laboratory measures, and lays foundation for better stratification of patients.

## Financial Disclosures of all authors (for the preceding 12 months)

**Chiara Milanese**

Stock Ownership in medically-related fields: None

Intellectual Property Rights: None

Consultancies: None

Expert Testimony: None

Advisory Boards: None

Employment: Erasmus University Medical Center Rotterdam, the Netherlands

Partnerships: None

Contracts: None

Honoraria: None

Royalties: None

Grants: None

Other: None

**Cesar Payan-Gomez**

Stock Ownership in medically-related fields:

None Intellectual Property Rights: None

Consultancies: None

Expert Testimony: None

Advisory Boards: None

Employment: Facultad de Ciencias Naturales y Matemáticas, Universidad del Rosario, Carrera 24, 63C-69 Bogotá, Colombia

Partnerships: None

Contracts: None

Honoraria: None

Royalties: None Grants: None

Other: None

**Marta Galvani**

Stock Ownership in medically-related fields: None

Intellectual Property Rights: None

Consultancies: None

Expert Testimony: None

Advisory Boards: None

Employment: University of Pavia, Pavia, Italy

Partnerships: None

Contracts: None

Honoraria: None

Royalties: None

Grants: None

Other: None

**Nicolás Molano González**

Stock Ownership in medically-related fields: None

Intellectual Property Rights: None

Consultancies: None

Expert Testimony: None

Advisory Boards: None

Employment: Facultad de Ciencias Naturales y Matemáticas, Universidad del Rosario, Carrera 24, 63C-69 Bogotá, Colombia

Partnerships: None

Contracts: None

Honoraria: None

Royalties: None

Grants: None

Other: None

**Maria Tresini**

Stock Ownership in medically-related fields: None Intellectual Property Rights: None

Consultancies: None

Expert Testimony: None

Advisory Boards: None

Employment: Erasmus University Medical Center

Rotterdam, the Netherlands Partnerships: None

Contracts: None

Honoraria: None

Royalties: None

Grants: Worldwide

Cancer Research Other: None

**Soraya Nait Abdellah**

Stock Ownership in medically-related fields: None

Intellectual Property Rights: None

Consultancies: None

Expert Testimony: None

Advisory Boards: None

Employment: None

Partnerships: None

Contracts: None

Honoraria: None

Royalties: None

Grants: None

Other: None

**Willeke M.C. van Roon-Mom**

Stock Ownership in medically-related fields: None

Intellectual Property Rights: None

Consultancies: None

Expert Testimony: None Advisory Boards: None

Employment: Leiden University Medical Center Partnerships: None

Contracts: None Honoraria: None Royalties: None Grants: None Other: None

**Silvia Figini**

Stock Ownership in medically-related fields: None

Intellectual Property Rights: None

Consultancies: None Expert Testimony: None Advisory Boards: None

Employment: University of Pavia, Pavia, Italy Partnerships: None

Contracts: None Honoraria: None Royalties: None Grants: None Other: None

**Johan Marinus**

Stock Ownership in medically-related fields: None Intellectual Property Rights: None

Consultancies: None Expert Testimony: None Advisory Boards: None

Employment: Leiden University Medical Center

Partnerships: None Contracts: None Honoraria: None Royalties: None

Grants: Alkemade-Keuls Foundation, Stichting Parkinson Fonds, Parkinson Vereniging, the Netherlands Organisation for Health Research and Development

Other: None

**Jacobus J. van Hilten**

Stock Ownership in medically-related fields: None

Intellectual Property Rights: None

Consultancies: None

Expert Testimony: None

Advisory Boards: None

Employment: Leiden University Medical Center

Partnerships: None

Contracts: None

Honoraria: None

Royalties: None

Grants: Alkemade-Keuls Foundation, Stichting Parkinson Fonds, Parkinson Vereniging, the Netherlands Organisation for Health Research and Development

Other: None

**Pier G. Mastroberardino**

Stock Ownership in medically-related fields: None

Intellectual Property

Rights: None

Consultancies: Neurological Institute “Casimiro Mondino” (non-profit)

Expert Testimony: None

Advisory Boards: None

Employment: Erasmus University Medical Center Rotterdam, the Netherlands

Partnerships: None

Contracts: None

Honoraria: None Royalties: None

Grants: AIFA (Italian Medicines Agency)

Other: Editorial Board member of *Neurobiology of Disease* and *Cell Death and Disease*; Associate Editor for *Frontiers in Cellular Neuroscience*

## Authors’ Roles

Chiara Milanese: research project execution, data analysis, manuscript review and critique

Cesar Payan-Gomez: statistical analysis design and execution, manuscript review and critique

Marta Galvani: statistical analysis execution

Nicolás Molano González: statistical analysis design and execution, manuscript review and critique

Maria Tresini: research project execution

Soraya Nait Abdellah: research project execution

Willeke M.C. van Roon-Mom: research project execution, manuscript review and critique

Silvia Figini: statistical analysis review and critique

Johan Marinus: statistical analysis and manuscript review and critique

Jacobus J. van Hilten: provided crucial intellectual input on study design; manuscript review and critique

Pier G. Mastroberardino: research project conception, organization, and supervision; data analysis; wrote the manuscript

